# Combined inhibition of SHP2 overcomes adaptive resistance to type 1 BRAF inhibitors in BRAF V600E-driven high-grade glioma

**DOI:** 10.1101/2024.12.21.629454

**Authors:** Abiola A. Ayanlaja, Michael Chang, Kriti Lalwani, Maria Ioannou, Jiawan Wang, Shreya Jagtap, Yanbo Yang, Robyn D. Gartrell, Christine A. Pratilas, Karisa C. Schreck

## Abstract

*BRAF*-mutant gliomas can be therapeutically targeted with BRAF mutant-selective inhibitors, yet responses are often transient due to short-term adaptive or long-term treatment-emergent resistance. We hypothesized that vertical inhibition of multiple signaling nodes could improve the durability of BRAF inhibition and prevent or overcome adaptive resistance. Using human tissue samples, we identified frequent RAS pathway reactivation in gliomas resistant to BRAF inhibitors, suggesting a common escape mechanism. Using patient-derived cell lines, we observed that upregulation of RAS activity was an adaptive response to BRAFi and that knockdown of SHP2, a central regulator of RAS activity, resulted in enhanced sensitivity to BRAF or MEK inhibition. Moreover, combined small molecule inhibition with SHP2 and BRAF or MEK inhibitors increased the depth and durability of ERK pathway inhibition, as well as prevented paradoxical upregulation of RAS activity. RNA sequencing analysis revealed deeper suppression of ERK transcriptional output with combined therapy, along with decreased reactivation of EGFR. Combined SHP2/BRAF small molecule inhibitors prevented growth and induced cell death in some cell line models. In cell lines with treatment-emergent resistance, moreover, combined SHP2 and BRAF inhibition overcame resistance to BRAF inhibitor monotherapy. *In vivo* orthotopic and patient-derived xenograft models confirmed enhanced tumor growth inhibition with combined therapy. Together, our findings demonstrate the critical role of RAS/ERK signaling reactivation in driving resistance to BRAF inhibition in glioma, and demonstrate the potential utility for adding SHP2 inhibitors to overcome resistance in *BRAF* V600E mutant glioma.

**Significance:** The addition of a SHP2i to BRAFi in BRAF-V600E glioma cells prevents tumor growth and can overcome resistance to BRAFi in preclinical models in vitro and in vivo.

## Introduction

Gliomas have increasingly been subcategorized based on their molecular alterations as formulated in the WHO 2021 central nervous system (CNS) integrated diagnostic criteria (1). Recent breakthroughs for some molecularly-driven subgroups of glioma have led to FDA-approvals of small molecule inhibitors for *BRAF*-mutant gliomas (2–6). While a large proportion of patients with *BRAF*-mutant glioma respond to targeted therapy, some do not, likely due to co-existing mutations that drive intrinsic resistance or due to tumor heterogeneity (5). In those for whom there is a clinical benefit, tumor progression may emerge after a period of response (2, 6). Treatment-emergent resistance can occur through adaptive mechanisms, by which upstream signaling components become activated due to release of negative feedback, as has been characterized initially in melanoma and confirmed in glioma (7–10). Alternatively, acquired resistance can result from the selective growth or survival advantage conferred by novel genomic alterations, as has been observed in gliomas (9, 11, 12). Co-occurring alterations in RAS family members (e.g., loss of *NF1)*, or loss of *CDKN2A/B* in pre-treatment tissue decrease likelihood of sustained clinical benefit from BRAFi targeted therapy, but other predictors of response are currently lacking (2, 5, 13).

Preventing or overcoming therapeutic resistance is an active area of investigation. In some cancers, combination therapies inhibiting additional vertical pathway components have proven effective at overcoming intrinsic resistance to BRAF inhibition, such as the addition of EGFR inhibition to BRAF/MEK inhibition for *BRAF* V600E colorectal cancer (14). Another approach is to inhibit independent pathways promoting survival, for example the addition of Hsp90 or CDK4/6 inhibitors to BRAF inhibitors (15, 16). Still another approach is to prevent adaptive resistance through autophagy, a described mechanism of escape for glioma (17). Given the paucity of other therapies for glioma and the relative success of BRAF inhibitors in this population, the need to understand adaptive and acquired resistance is particularly urgent.

Previously, we and others demonstrated that acquired resistance in *BRAF*-altered gliomas occurs through co-mutations that preserve ERK signaling overactivity (9, 11). Here, we investigate treatment-emergent resistance at a protein level in human tissue samples, confirming ERK-dependence upon resistance. This finding raised the appeal of a vertical inhibition strategy. One attractive target for combination is the SH2 domain-containing tyrosine phosphatase 2 (SHP2), encoded by the gene *PTPN11,* which is a ubiquitously expressed non-receptor protein tyrosine phosphatase. Due to its critical role in signal transduction between receptor tyrosine kinases (RTKs) and RAS, SHP2 modulates RAS-dependent ERK, mTOR, and JAK/STAT signaling pathways (18, 19). SHP2-dependent RAS activation can drive adaptive resistance to MEK inhibitors in some cancers, and SHP2 inhibition with small molecule inhibitors (SHP2i) can prevent or overcome adaptive resistance to MEK inhibitors (20–23).

We evaluated adaptive and acquired resistance in *BRAF*-mutant glioma models and examine the potential efficacy of a SHP2 inhibitor-based vertical combination strategy. We compare the efficacy of combined small molecule inhibitor approaches to prevent adaptive resistance or overcome acquired resistance to BRAF or MEK inhibitors *in vitro* and *in vivo*. Our findings provide the foundation for a clinical strategy utilizing SHP2 inhibitors in the care of patients with *BRAF* V600E altered cancers.

## Materials and Methods

### Human specimens

Human tissue specimens are previously described in a prior study and were obtained in compliance with Institutional Review Board (IRB) regulations at Johns Hopkins Hospital (Baltimore, MD; IRB00158788) and elsewhere as described when they were first published (11).

### Cell lines

Five *BRAF* V600E mutant glioma cell lines were maintained in 5% CO_2_ at 37 °C as previously described (11). DBTRG-5MG cell line (DBTRG, RRID: CVCL_1169), obtained from ATCC, was grown in RPMI1640 supplemented with 10% fetal bovine serum (FBS) and 1% penicillin/streptomycin. NMCG1 (RRID: CVCL_1608) and NGT41 were grown in DMEM supplemented with 10% FBS and 1% penicillin/streptomycin (24). B76 and MAF794 were grown in Opti-MEM Reduced Serum Media supplemented with 15% FBS and 1% penicillin/streptomycin. BRAFi-resistant DBTRG, NGT41, and B76 lines were generated via chronic exposure to increasing doses of dabrafenib monotherapy starting at 30 nM and increasing to 10 µM. MAF749 parental and resistant to vemurafenib were a gift from Dr. Jean Mulcahy Levy, generated as previously described (17). Cells were maintained in 10 µM dabrafenib or vemurafenib every passage for 90 days to ensure stability. For resistant cell lines cultured in the presence of chronic BRAFi, targeted therapy was withdrawn for 3 days prior to experiments.

### Cell Growth Assays

All cell lines were stably transduced with Incucyte Nuclight Red Lentivirus (EF1α, puro, Sartorius #4476) and selected in 2 µg/ml puromycin for stable expression of nuclear-restricted red fluorescent label. For growth assays, cells were plated in a 96-well plate at a density of 1.0-1.5 × 10^3^ cells/well and treated with drug 24 hours after plating. Cell growth was assessed with the IncuCyte^TM^ S3 Live-Cell Imaging System tracking area of red nuclei or monitoring confluence area. Both measures are normalized to day zero for that specific frame to control for differences in plating density. Where appropriate, synergy scores were calculated using the web based “SynergyFinder” to analyze efficacy of pairwise combination therapy (25). Using the Loewe’s model, a synergistic score was determined as values ≥ 10 while additive interaction is between 0 and 10.

### Immunoblotting

Cells were collected and prepared as previously described (26). Antibodies against RAS (ab52939) was purchased from Abcam. Antibodies against NRAS, HRAS, ARAF, CRAF, GAPDH, SHP2, total ERK, phospho-MEK, phospho-ERK, phospho-AKT(Ser473), phospho-AKT(Thr308), and phospho-EGFR were from Cell Signaling.

### *PTPN11* Knockdown (siRNA)

*PTPN11* knockdown was performed using SMARTPool ON-TARGETplus human *PTPN11* siRNA (#L-003947–00-0005, Dharmacon), with non-targeting pool (#D-001810–10-05, Dharmacon) as a control, to knockdown (KD) the expression of *PTPN11* or non-targeting control (NTC), respectively. DharmaFECT 1 Transfection Reagent (#T-2001–02, Dharmacon) was used to facilitate transfection according to manufacturer protocols as previously described (21). The knockdown of SHP2 was confirmed using immunoblot for SHP2 protein.

### RNA Sequencing

B76, DBTRG, NGT41, and MAF794 cell lines were cultured and treated in triplicate with either DMSO, dabrafenib (100 nM), TNO155 (3 µM), or a combination of dabrafenib and TNO155. For transcriptome sequencing, libraries were prepared following manufacturer protocol using the TruSeq Stranded Total RNA LT Sample Prep Kit and run on the Illumina HiSeqX platform to generate paired-end reads of 150 base pairs. Illumina’s CASAVA 1.8.4 was used to convert BCL files to FASTQ. Data were further processed in R version 4.3.1. RSEM v1.3.0 was used to align reads to the hg38 reference genome and quantify transcript expression (27). Differential expression analysis was performed with DESeq2 v1.40.2 (28). Subsequent gene set enrichment analysis (GSEA) was performed with the clusterProfiler v4.8.3 package in R (29). After pre-ranking all genes by signed log2 fold change*-log10(p-value) calculated from DESeq2, we utilized the MsigDBR v7.5.1 package to query gene sets from the Hallmark (H), KEGG (CP:KEGG), and chemical and genetic perturbations (CGP) collections in our analysis (30, 31). We also conducted hypergeometric enrichment analysis with the ClusterProfiler package. Hallmark gene sets and oncogenic signature gene sets from the Molecular Signatures Database were used for pathway annotation (30). Top 5 pathways ranked by adjusted pval were selected. We then performed differential expression analysis comparing combined BRAFi with SHP2i to BRAFi alone. GSEA analyses was performed with the ranked gene list and results were visualized using GSEA running enrichment plot using enrichplot v1.24.4 package (29).

### Active RAS pull down assay

Cells were washed and collected for GTP-bound RAS quantification using the Cell Signaling active RAS detection kit (# 8821) or lysed in NP40 lysis buffer for total protein expression. Protein was quantified using the Pierce BCA protein assay kit (# 23227, Thermo Fisher Scientific). Equal amounts of protein were separated via SDS-PAGE, transferred onto nitrocellulose membranes, immunoblotted with target protein’s primary and secondary antibodies. Protein expression was detected using chemiluminescence with Immobilon western chemiluminescent HRP substrate (#WBKLS0500, Millipore). Imaging was performed using a ChemiDoc touch imaging system (Bio-Rad).

### *In vivo* mouse studies

6-8 weeks old NOD SCID gamma (NSG, # 005557, Jackson Laboratories) female or male mice were implanted subcutaneously with cell lines or PDX tumor fragments. When heterotopic xenografts reached approximately 400 mm^3^ in size, mice were randomized into treatment groups by an R-generated algorithm that groups animals into best-case distribution with similar tumor burden and standard deviation in each treatment group. Vehicle, dabrafenib (15 mg/kg, Selleckchem; dissolved in 0.2% Tween 80, 0.5% hydroxypropyl methyl cellulose, pH 8.0), and TNO155 (7.5 mg/kg, Novartis; dissolved in 0.1% Tween 80 and 0.5% methyl cellulose), or their combination, were administered twice daily. Drugs were administered via oral gavage for up to 28 days and mice were carefully monitored for symptoms and sacrificed when tumors reached protocol limits. Tumor volume was calculated as previously described (32).

For orthotopic intracranial implantation, 500,000 cells were injected intracranially into the right striatum of 6-8 weeks old immune compromised NSG mice as previously described (26). 5 days after injection, mice were randomly distributed into four treatment groups. 50 mice were grouped in a cohort of 4 groups, including 10 mice per arm of vehicle, TNO155, dabrafenib, and the combination. Drugs were administered twice daily via oral gavage starting 7 days after implantation (5 days on / 2 days off) for six weeks. Mice were monitored daily by an independent, blinded evaluator and sacrificed with the development of neurological symptoms. Kaplan-Meier survival curves were plotted. Animal survival difference was validated using the log-rank test in Graph Prism v10.2 software (GraphPad software, Inc.). All murine experiments were approved by the Institutional Animal Care and Use Committee at Johns Hopkins under protocol # MO21M387.

### Statistical Analyses

Differences in cell growth was calculated using mean and standard error. Unpaired t-test was used to determine differences between normally distributed conditions. Wald test was used alongside DESeq to identify differentially expressed genes and visualized via volcano plot with thresholds for DEGs defined as Log2-Fold-change ≥ 1 OR ≤ −1 (vertical dashed lines) and Benjamini-Hochberg corrected p-values ≤ 0.05 (horizontal dashed line). Student’s t-test was used to test for differences in transcriptomic signatures among cell lines treated with different conditions. Adjusted p-values from gene set enrichment analysis were calculated using the Benjamini-Hochberg method (default).

## Results

### RAS reactivation is associated with resistance to ERK signaling inhibition in human glioma

In order to understand mechanisms of adaptive resistance to BRAFi/MEKi in glioma, we evaluated protein expression in a cohort of human *BRAF* V600E mutant glioma specimens obtained preceding treatment with BRAFi/MEKi and compared them against tissue obtained at the time of progression (**Figure 1A**). Of note, all the patients from whom pre-treatment samples were collected and included in this analysis, experienced initial response to BRAFi/MEKi (11). Tumor specimens were evaluated for ERK signaling activity and for expression of phospho-EGFR and PDGFRβ, given their high frequency of dysregulation in glioblastoma (33). Each post-treatment sample demonstrated preserved MEK/ERK signaling at progression, along with a trend in the cohort towards increased PDGFRβ expression and EGFR activation (**Fig 1A-C**). In two individual patients with paired tumor samples (**Fig 1C**), immunoblot analysis revealed an increase in pEGFR and PDGFRβ in both post-versus pre-BRAFi pairs. These results support prior findings that cancers with *BRAF* V600E are addicted to ERK signaling and mechanisms that drive resistance converge upon its reactivation (7, 34).

**Figure 1:**
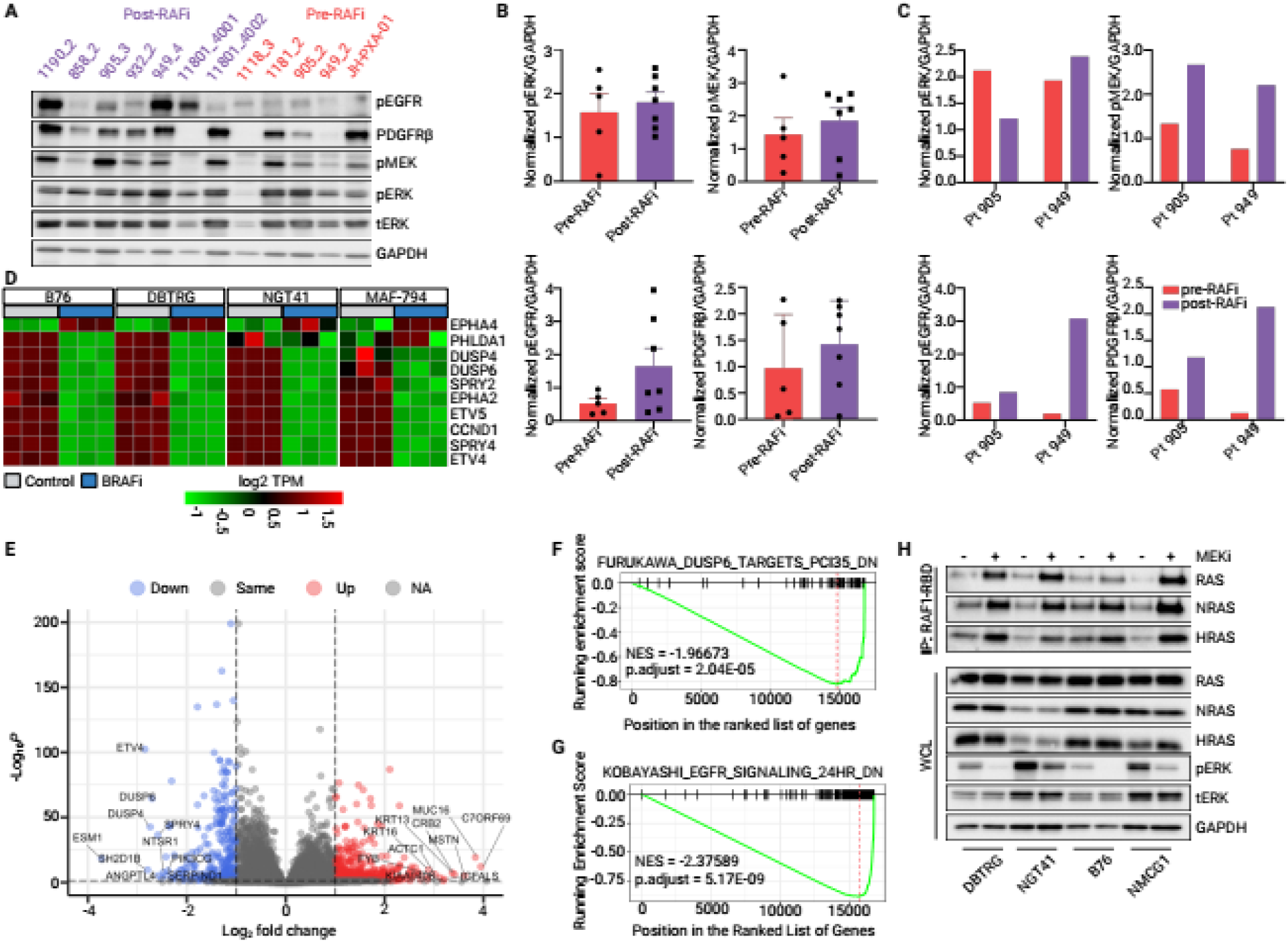
RAS reactivation is a common response to BRAF inhibition in gliomas. A) Immunoblot of *BRAF* V600E glioma samples from human patients before (pre-RAFi) or after (post-RAFi) RAF inhibitor treatment B) quantified and normalized to GAPDH across all samples or quantified for paired, same-patient samples (C). D) MAPK Pathway Activity Score (MPAS) genes in four *BRAF* V600E mutant glioma lines before or after treatment with BRAFi (dabrafenib,100nM) for 24 hours in triplicate. E) Volcano plot labeling top 10 significant positive and negative LFC (log fold change) transcripts in BRAFi-treated (dabrafenib, 100nM)-treated cell lines compared to controls. Thresholds for DEGs defined as Log2-Fold-change ≥ 1 OR ≤ −1 (vertical dashed lines) and Benjamini-Hochberg corrected p-values ≤ 0.05 (horizontal dashed line). F, G) Genset enrichment analysis pooled across all lines post-versus pre-treatment for two gene sets of interest. H) RAS-GTP was assayed in four *BRAF* V600E mutant glioma cells treated with MEK inhibitor (trametinib, 100nM) for 24 hours using the RAF1-binding assay. WCL = whole cell lysate.

### *BRAF* V600E glioma cells display heterogenous BRAFi/MEKi sensitivity

To understand the biological relevance of *BRAF-*mutant cell lines for evaluating resistance, we sought to clarify the dependence of RAF-ERK signaling on oncogenic *BRAF* V600E versus exogenous growth factors. We assessed signaling as measured by phospho-ERK levels on immunoblot in the presence or absence of 10% FBS and observed two different patterns of response. In some *BRAF* V600E mutant cell lines (DBTRG, B76, NGT41, and MAF794), we observed ERK activity is independent of upstream RTK signaling as evidenced by no signaling change in the presence or absence of serum (**Fig S1A-D**). In another line (NMCG1), we observed serum-dependence of ERK and PI3K signaling (pERK, pS6, pAKT), indicating relatively less dependence on *BRAF* V600E. In addition to BRAF mutation, NMCG1 had MET, EGFR, and BRAF amplification, which are likely mechanism of resistance to RAFi (35), (**Fig S1E**). The dependence on growth signaling upstream of RAS was inversely related to BRAFi/MEKi sensitivity (**Fig S1F-K**).

We next examined whether short-term adaptive signaling changes in response to BRAFi treatment would alter transcriptional output in a manner that correlated with growth inhibition to BRAFi. We analyzed differential gene expression in four sensitive *BRAF* V600E cell lines before or after BRAFi treatment using bulk RNAseq. We evaluated transcriptional signatures correlating with MEKi sensitivity or MEK-ERK functional dependence previously validated in a range of solid cancers (7, 36). We also calculated the MAPK pathway activity score (MPAS), a 10-gene transcriptional output score derived as a consensus signature from several overlapping ERK output/ MEK inhibitor sensitivity scores, which correlates with sensitivity to MAPK pathway inhibitors (37). MPAS gene expression showed similar inhibition of ERK output in four *BRAF* V600E cell lines in response to BRAFi (**Fig 1D**). Evaluation of another validated MEK-ERK functional output signature showed similar inhibition of ERK output across all cell lines (**Fig S2A**) (36).

To determine overall transcriptional changes in response to short term BRAF inhibition, we identified the top ten up- and down-regulated genes using a consensus analysis of all four cell lines. We observed consistent downregulation of ERK signaling related genes and downstream markers of ERK pathway activity including *ETV4*, *DUSP6, DUSP4,* and *SPRY4* (**Fig 1E**, **Fig S2B-E**). Relatively fewer genes were upregulated in response to BRAFi. We also performed gene set enrichment analysis (GSEA) to identify the most enriched Hallmark pathways, as well as change in curated gene sets associated with ERK signaling. One of the most significantly enriched gene sets was one inhibited by DUSP6, indicative of profound pERK inhibition after BRAFi relative to control (Furukawa-Dusp6-targets-PCI35-down, p-adj < 0.001; **Fig 1F**). The most significantly enriched gene set following BRAFi was one downregulated with EGFR inhibition, suggesting EGFR signaling is upregulated with adaptive resistance (Kobayashi-EGFR-signaling-24hr-dn; NES = −2.37; p-adj < 5×10^-9^; **Fig 1G**). Genes upregulated with EGFR inhibition were also significantly upregulated in cells following BRAFi (NES = 1.59, p-adj 0.02; **Fig S2F**). Conversely, genes upregulated with KRAS activation were negatively enriched (Hallmark_KRAS_signaling_up; NES −1.68, p-adj = 0.001, **Fig S2G**), a similar trend as shown for the mTORC1-dependent gene signature (Hallmark_MTORC1_signaling; NES −1.64, p-adj 0.01; **Fig S2H**).

Studies in other cancers, as well as the human glioma specimens, have demonstrated that resistance to RAF inhibitors often occurs through reactivation of RAS-ERK signaling. To determine whether this was true in our glioma models, we quantified RAS-GTP levels in response to short-term MEKi versus control using a RAF1-RAS binding domain (RAF1-RBD) based immunoprecipitation to extract the fraction of RAS that is GTP-bound and able to bind RAF1. Treatment with MEKi (100 nM, trametinib) increased GTP-bound—but not total—RAS within 24 hours of treatment (**Fig 1H**), demonstrating that loss of feedback inhibition from ERK to RAS leads to reactivation of ERK signaling.

### SHP2 inhibition does not modulate ERK signaling or proliferation in *BRAF* V600E mutant gliomas

Given our observation that RAS upregulation in response to BRAFi/MEKi in glioma is associated with a rebound in pERK activity and contributes to limited response duration to BRAFi or MEKi, we postulated that inhibition of SHP2, a critical regulator of RAS activity, might prevent adaptive resistance. First, to investigate the role of SHP2 in *BRAF* V600E glioma, we transiently reduced SHP2 expression in *BRAF* V600E glioma cells using pooled siRNA targeting *PTPN11*. As expected, *PTPN11* knockdown alone did not significantly reduce ERK activity or cell viability and growth (**Fig 2A, B**), consistent with published reports that *BRAF* V600E drive signaling independently of upstream RAS activity (8).

**Figure 2:**
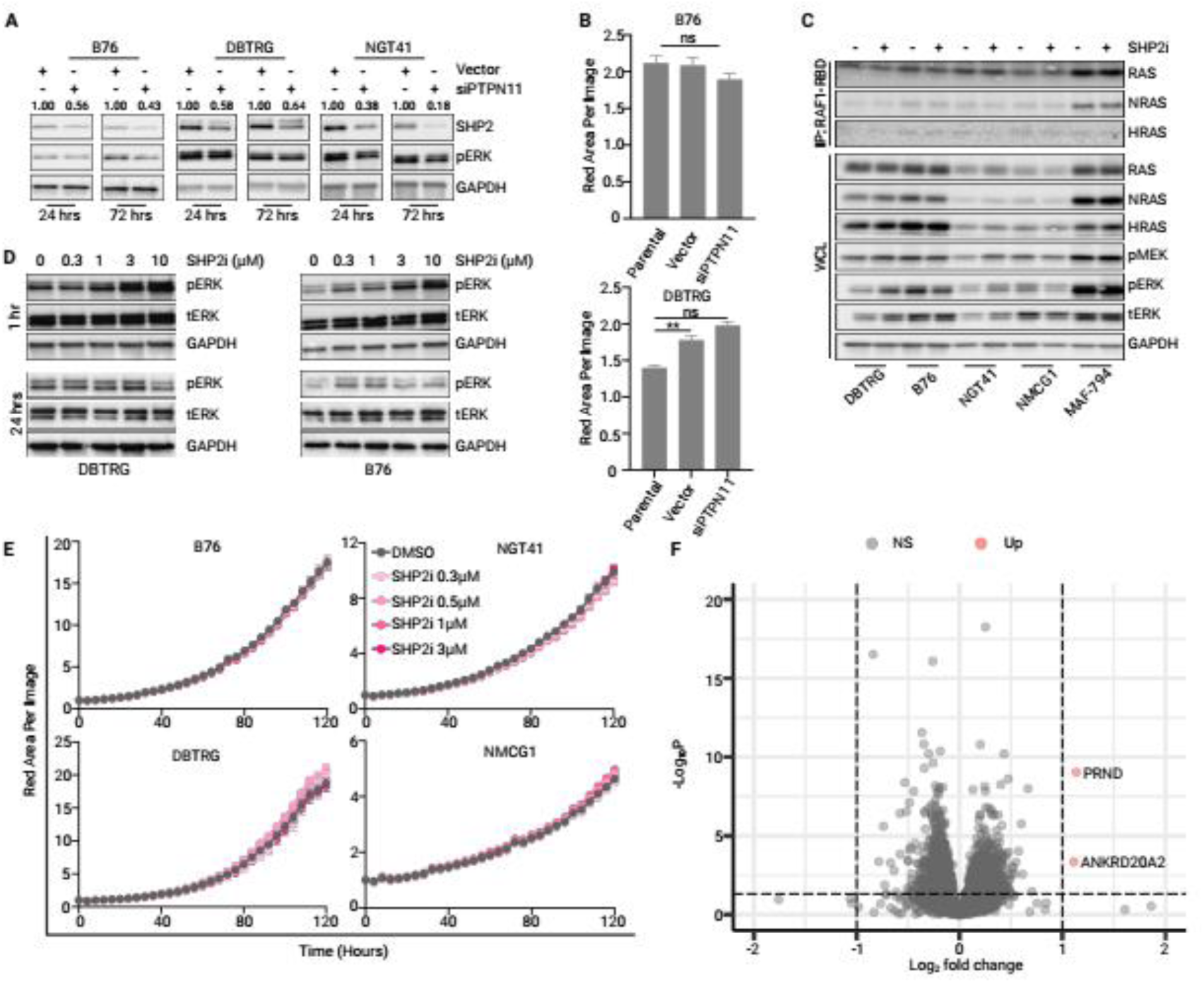
SHP2i monotherapy is ineffective at inhibiting ERK signaling or proliferation in *BRAF* V600E mutant gliomas. A) siRNA mediated knockdown of PTPN11 (siPTPN11) in three *BRAF* V600E mutant cell lines evaluated using immunoblot for SHP2 and pERK proteins at 24 and 72 hrs post transfection. Quantification of SHP2 relative to GAPDH shown above the appropriate band. B) Relative cell number measured by a red nuclear label (dsRed) 4.5 days after transduction with either vector or siPTPN11. C) RAS-GTP pulldown assay 24 hours after TNO155 treatment (3 µM); WCL = whole cell lysate. D) Two *BRAF* V600E mutant glioma lines (DBTRG and B76) were treated with increasing doses of TNO155 (0.3 µM-3 µM) at 1 hour and 24 hours after treatment. E) Cell growth over time following treatment with increasing doses of TNO155 (0.3-3 µM), normalized to baseline cell density per image as quantified by red nuclear area. F) Volcano plot of differentially expressed transcripts in response to TNO155 (3 µM) monotherapy averaged across 4 cell lines.

We next evaluated the impact of the small molecule SHP2i, TNO155, in *BRAF* V600E mutant glioma cell lines. We evaluated whether treatment with a SHP2i would impact the fraction of GTP-bound RAS across five different *BRAF* V600E mutant glioma lines and saw no change (**Fig 2C**). The SHP2i did not inhibit RAS or ERK activity, instead causing dose-dependent induction of pERK in some cell lines (**Fig 2D**). In one line with demonstrated dependence on upstream growth factors for pathway activity, NMCG1, pERK was partially inhibited by the SHP2i but growth was not impacted (**Fig 2C**). In fact, SHP2i treatment did not inhibit cell growth in any of the cell lines examined (**Fig 2E**). Pooled analysis of gene expression changes induced by SHP2i monotherapy across four cell lines revealed no discernible effect on ERK transcriptional output or other pathways relevant for cell signaling (**Fig 2F**, **S3A-D**).

### SHP2 inhibition augments MEKi by preventing compensatory RAS reactivation

Despite the absence of single agent activity of the SHP2i, we hypothesized that the combination of a SHP2i with MEKi would deepen ERK pathway inhibition by preventing compensatory reactivation of RAS. While *PTPN11* knockdown alone had no effect on cell growth, we observed that *PTPN11* knockdown combined with the MEKi trametinib enhanced growth inhibition compared to either condition alone (**Fig 3A**). We asked whether compensatory RAS reactivation would be inhibited, and found that the addition of SHP2i to MEKi prevented RAS reactivation at 24 and 48 hours as measured by levels of GTP bound RAS (**Fig 3B**), comparable to the effect seen in *NF1*-deficient cancer models (21).

**Figure 3:**
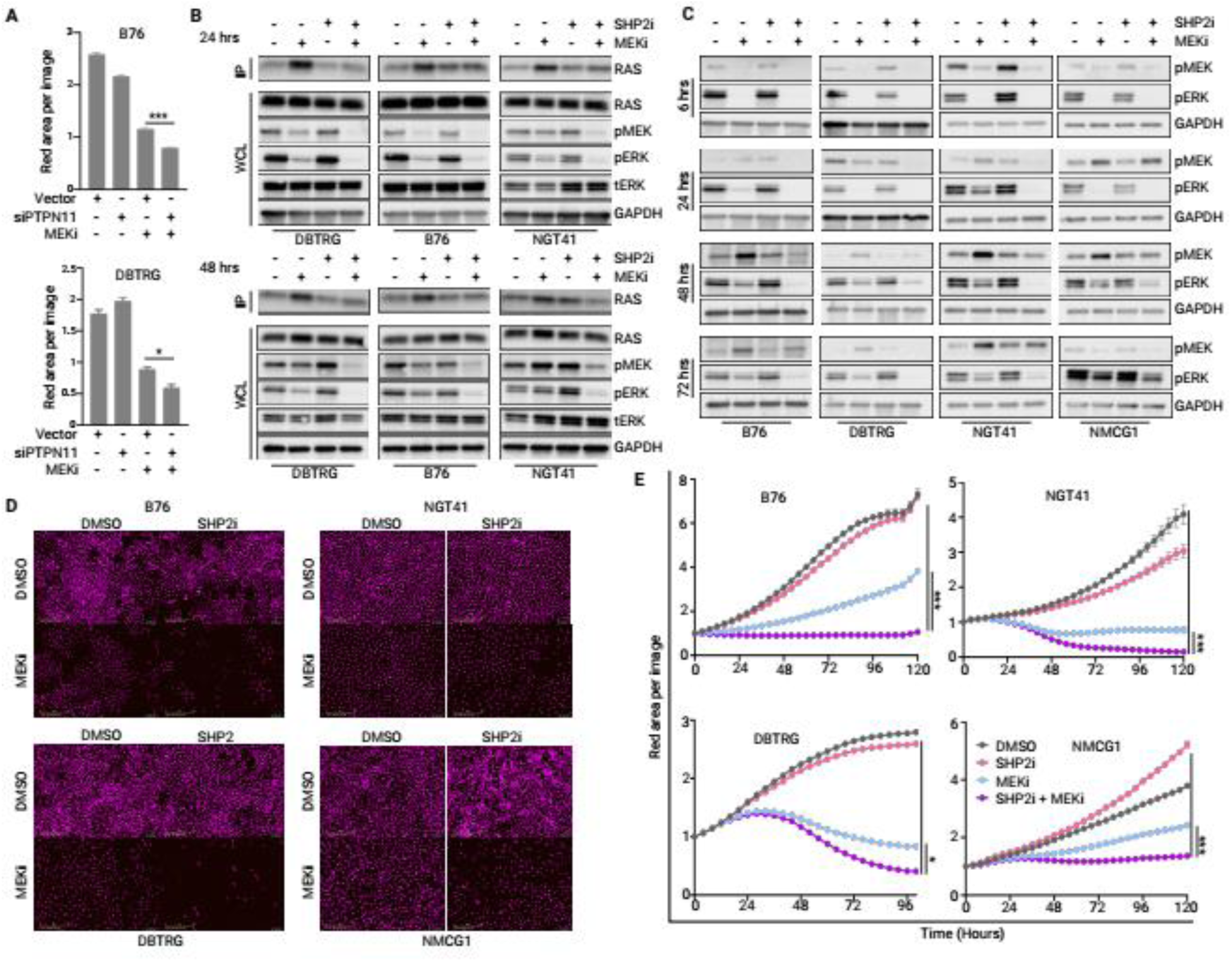
SHP2 inhibition augments MEKi by preventing compensatory RAS reactivation. A) B76 and DBTRG cells were transduced with either vector or siPTPN11, then treated with trametinib (30 nM) for 4.5 days. Cell growth was quantified as nuclear-red area per image. B) RAS activity was quantified using RAS-GTP pulldown assay in cell lines treated with DMSO, TNO155 (3 µM), trametinib (30 nM), or the combination at 24 or 48 hours after treatment. IP=IP: RAF1-RBD, WCL= whole cell lysate. C) Immunoblots of ERK pathway activity collected 6 - 72 hours following treatment with DMSO, TNO155 (3 µM), trametinib (30 nM), or the combination. D, E) Cell density, with nuclear fluorescence pseudo-colored magenta, following treatment with TNO155, trametinib or combination was quantified over time on 4x images using IncuCyte. Error bars indicate mean ± SEM (*p ≤0.05, *** p<0.0001).

We assessed the effect of SHP2i combined with MEKi on MEK-ERK signaling over time (**Fig 3C**). We observed profound pERK inhibition at short intervals with MEKi alone (6 hours), but reactivation after 24-48 hours. Notably, while SHP2i alone did not inhibit ERK activity, its addition to a MEKi increased the durability of ERK inhibition to as long as 72 hours. This finding was also true to a lesser extent in NMCG1, which was relatively insensitive to MEKi monotherapy. Consistent with the effect of combination therapy on pERK inhibition, the combination of MEKi with SHP2i inhibited cell growth more profoundly than either monotherapy in all *BRAF* V600E lines examined (**Fig 3D, E**).

### SHP2 inhibition augments BRAFi in glioma cells

While the addition of a SHP2i to MEKi augmented durability of ERK inhibition and growth inhibition in *BRAF* V600E glioma cell lines, concerns have been raised over the tolerability of this combination (38). In an effort to improve tolerability, we evaluated the effects of combining SHP2i and BRAF inhibitor, which may have an improved therapeutic index, due to the opposing effects on phospho-ERK in normal tissues. We treated glioma cells with the combination of TNO155 (3 µM) and dabrafenib (100 nM) and observed substantial growth inhibition in response to the combination (**Fig 4A, B**). We then asked whether augmenting BRAF inhibition with *PTPN11* knockdown would have a similar effect. We used pooled siRNA to knock down *PTPN11* alone or in combination with the BRAFi dabrafenib. We observed enhanced growth inhibition and decreased proliferation with *PTPN11* knockdown in addition to BRAFi (**Fig 4C**). Upon evaluation of combinatorial effect size, we found that dabrafenib and TNO155 induced a synergistic effect on all evaluated *BRAF* V600E mutant cell lines (**Fig 4D, S4A-C**). The combination of BRAFi and SHP2i resulted in sustained suppression of ERK signaling (pERK levels) for up to 72 hours (**Fig 4E**), similar to the effect seen with MEKi, supporting improved durability of ERK signaling inhibition with a combined vertical inhibition strategy. Notably, the cell line with intrinsic resistance to BRAFi, NMCG1, demonstrated partial pERK inhibition and growth suppression with the addition of SHP2i, likely through inhibition of upstream inputs to ERK signaling (**Fig 4B, E**).

**Figure 4:**
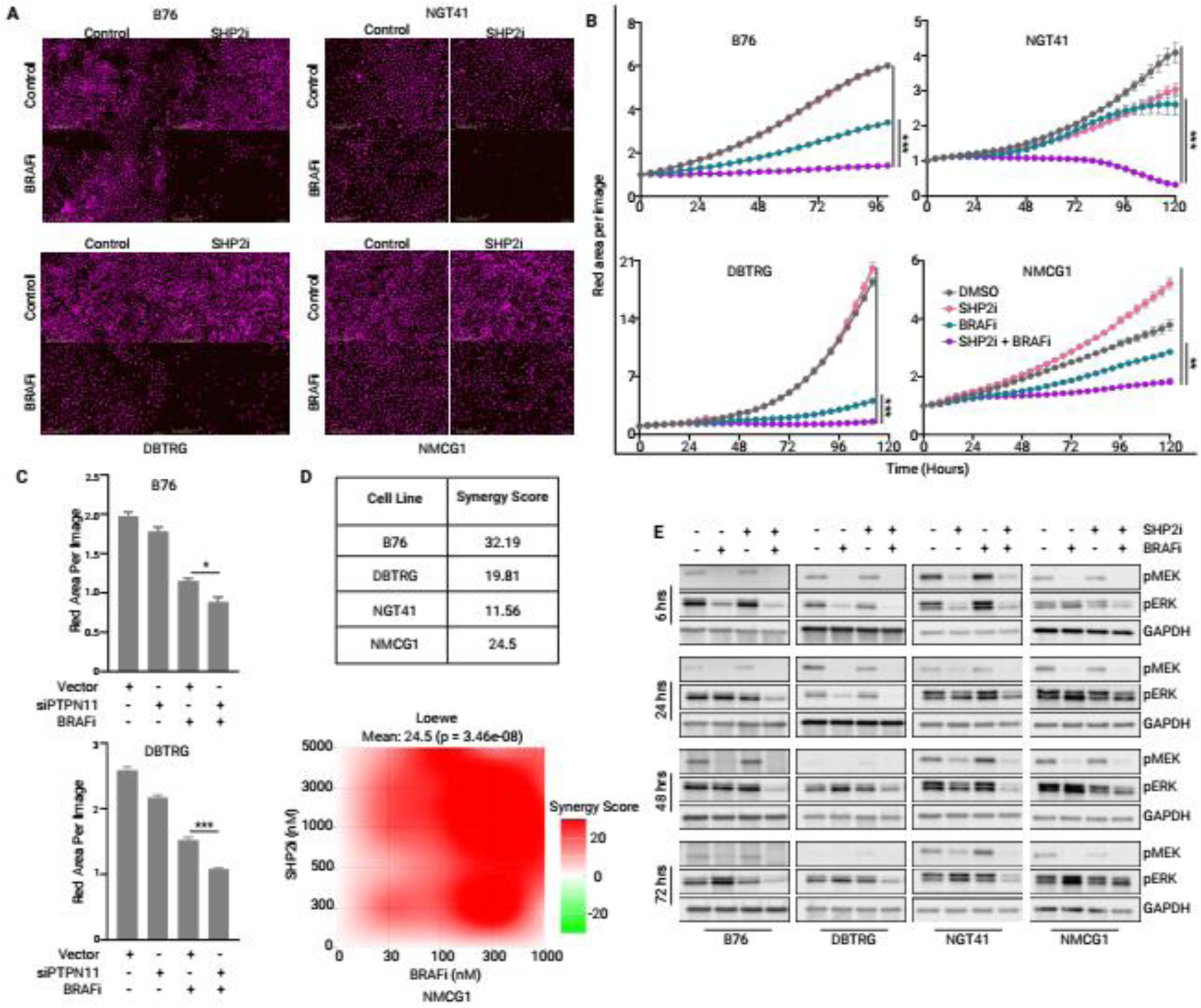
SHP2 inhibition augments BRAFi in glioma cells. A/B) Representative images and growth curves of cells treated with vehicle, TNO155 (3 µM), dabrafenib (100 nM), or the combination as assessed by live-cell imager. Magenta indicates nuclear red fluorescence label. C) Growth of B76 and DBTRG cells transduced with either empty vector or siPTPN11, then treated with dabrafenib (100nM) for 4.5 days quantified as nuclear-red area per image. Error bars indicate mean ± SEM (*p ≤0.05, ***p ≤0.0001. D) Synergy scores for each cell line were calculated using the Loewe’s model. Additive - Score < 10, synergistic - Score ≥ 10. The bottom panel shows representative image of synergy heatmap in NMCG1 cell line. E) Immunoblots of ERK pathway activity collected 6 - 72 hours following treatment with DMSO, TNO155 (3 µM), dabrafenib (100 nM), or the combination.

We sought to understand the effect of combined BRAFi with SHP2i on cell death. The combination of SHP2i with MEKi showed induction of cleaved PARP at 48-72 hours after treatment in sensitive lines (**Fig S4D-F**). Similarly, the combination of SHP2i with MEKi showed induction of cleaved PARP 48-72hrs following treatment, but BRAFi sensitive cell lines (DBTRG and NGT41) showed induction of cleaved-PARP as early as 6 hours after treatment (**Fig S4G-J**).

Given the concerns about the clinical tolerability of MEKi/SHP2i combinations, we measured the effect of BRAFi/SHP2i on human astrocytes *in vitro*. We observed no toxic effects of combined dabrafenib with TNO155 on normal human astrocytes at the dose regimen tested, with normal cell growth compared to controls (**Fig S4H**).

### Vertical inhibition increases RAS-ERK pathway suppression

Based on the findings above, we hypothesized that combined inhibition with SHP2i and BRAFi would prevent resistance to targeted therapy by suppressing RAS-ERK signaling more deeply than monotherapy and preventing compensatory upregulation of parallel pathways. To evaluate this hypothesis, we used bulk RNAseq to identify transcriptional changes and pathway-level modulation associated with BRAFi and SHP2i treatment. We compared the differentially expressed genes (DEGs) for each monotherapy or the combination against control across all cell lines (**Fig 5A**). More genes were differentially expressed with combined therapy than with either monotherapy and a large subset of DEGs overlapped between BRAFi monotherapy and the combination. In contrast, we observed very few DEGs in response to SHP2i monotherapy, and none overlapped with the combination (**Fig 5A**).

**Figure 5.**
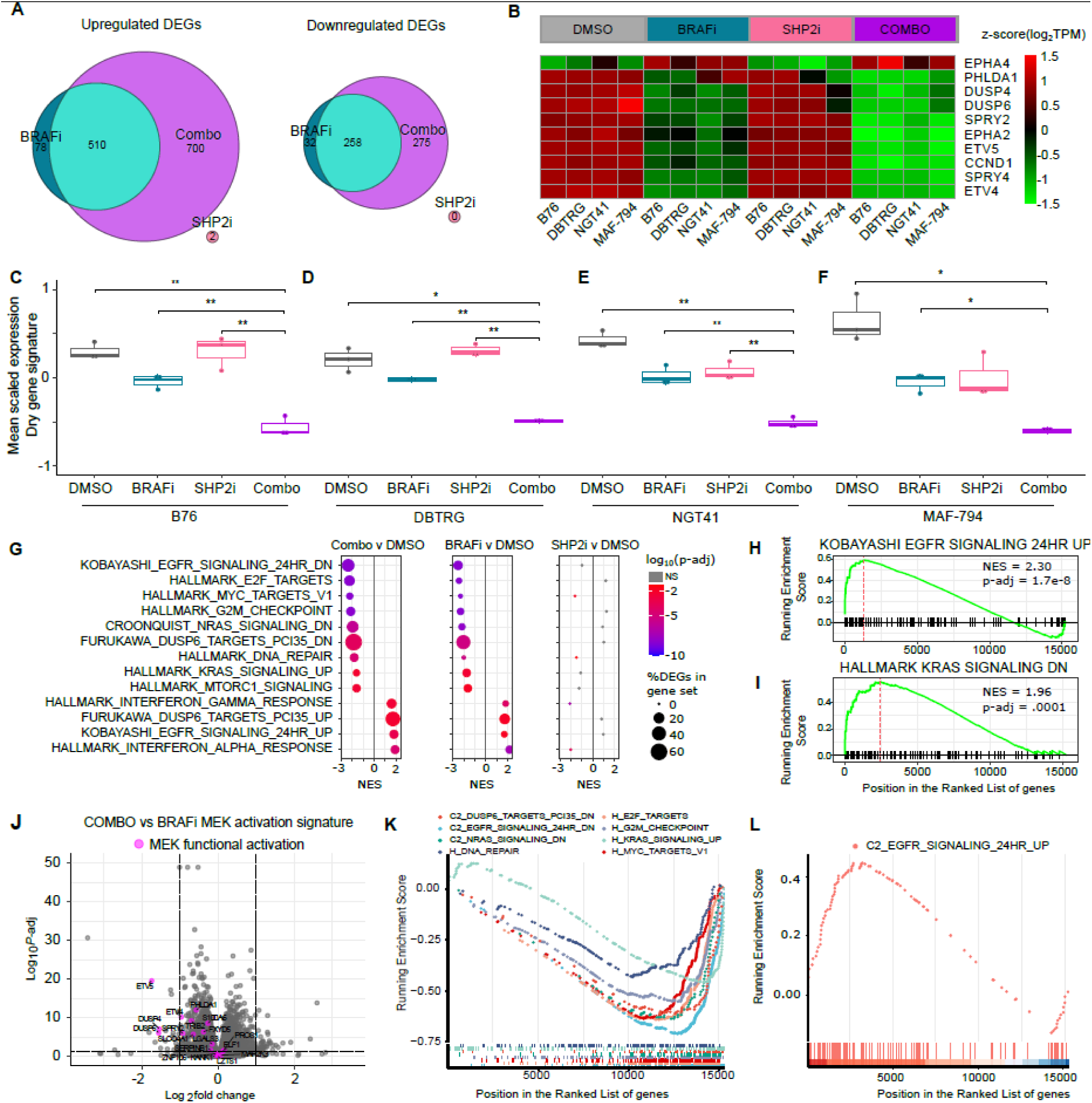
Combined therapy increases ERK pathway suppression in human glioma cell lines. A) Venn diagrams displays counts of significantly upregulated and downregulated genes common between dabrafenib (BRAFi, 100 nM), TNO155 (SHP2i, 3 µM) and combination (Combo) groups compared with DMSO-treated control across four BRAF-mutant glioma cell lines (B76, DBTRG, NGT41, MAF-794). Thresholds for DEGs were defined as Log2-Fold-change ≥ 1 OR ≤ −1 and p-adjusted ≤ 0.05. B) Heatmap of log_2_TPM of the 10 genes in MPAS gene signature in four BRAF-mutant lines across treatment conditions. C-F) Box-and-whisker plots display mean scaled expression of an 18-gene signature for MAPK addiction per condition. G) Dot plots display normalized enrichment scores and adjusted p-values from GSEA compared against the control condition. The top 8 Hallmark gene sets (from Combo vs DMSO comparison) with lowest adjusted p-value plus five additional hand-curated gene sets of interest for MAPK signaling are displayed. H/I) Genset enrichment analysis pooled across all lines for the combination versus control for two gene sets of interest. J) Volcano plots highlight the change in MAPK-associated genes with combination versus dabrafenib. K) GSEA multi-sample running enrichment plot comparing combination versus dabrafenib for pathways uniquely downregulated or L) upregulated.

We analyzed MAPK pathway activity using the MPAS gene expression score. As anticipated, it was unaffected by SHP2i treatment compared to controls, but the addition of SHP2i to BRAFi treatment further suppressed the MPAS score over BRAFi alone (**Fig 5B, S5A-D**). We then examined the mean scaled expression of MAPK related genes associated with MEK addiction and response to MEKi (36). For all cell lines, we observed modest inhibition of genes in the signature with BRAFi monotherapy, but profound inhibition with the addition of SHP2i (**Figure 5C-F**). Of note, SHP2i alone had no effect on two cell lines (**Fig 5C-D**), but resulted in partial inhibition in the other two lines (**Fig 5E, F**). However, SHP2i monotherapy had no effect on MPAS signature in all cell lines, suggesting increased signature specificity in these glioma models (**Fig S5E-H**).

To investigate other affected pathways in an unbiased fashion, we identified top differentially expressed pathways using gene set enrichment analysis (GSEA) performed using the Hallmark, KEGG, and CGP gene set collections (**Fig 5G**). We identified the top eight most enriched Hallmark pathways for the dabrafenib monotherapy condition against control, as well as five gene sets of interest identified as differentially regulated with BRAFi monotherapy. Among the most significantly enriched gene sets were those related to cell growth and cell cycle regulation, interferon α/γ response, and signaling downstream of DUSP6, EGFR, and RAS (**Figure 5G-I**). SHP2i monotherapy did not significantly impact any MAPK-associated pathways (**Fig 5G**). Notably, combined therapy impacted the same pathways as BRAFi, which were associated with cell cycle progression, growth, and RAS signaling. Compared with BRAFi alone, MEK functional activation signature genes were more profoundly suppressed (**Fig 5J; Fig S5I-L**). Other affected pathways were also more significantly up or down regulated with the combination than with BRAFi monotherapy, speaking to the amplified effect of combined vertical pathway inhibition (**Fig 5K-L**).

### Combination of BRAFi and SHP2i is effective in BRAFi-resistant glioma cells

Treatment-emergent resistance to BRAFi/MEKi is well documented in *BRAF* V600E mutant cancers (2, 5, 39, 40). To assess whether SHP2i can restore sensitivity in BRAFi-resistant glioma lines, we generated dabrafenib-resistant *BRAF* V600E altered glioma cell lines through prolonged culture in increasing doses of dabrafenib. Acquired dabrafenib resistance was confirmed by evaluating growth inhibition to dabrafenib between resistant and parental lines (**Fig 6A, B**). We also examined the vemurafenib-resistant cell line, MAF-794, demonstrating its growth insensitivity to vemurafenib (**Fig 6C**). We compared the effect of dabrafenib on ERK phosphorylation in parental/ resistant pairs (**Fig 6D, E**). While many cell lines insensitive to dabrafenib demonstrate transiently decreased ERK phosphorylation in the presence of dabrafenib (41), dabrafenib-resistant glioma lines demonstrated reduced inhibition of pERK at 1 hour, suggesting an adaptive means of maintaining ERK activity. The vemurafenib-resistant line, however, exhibited no reduction in ERK activity at either the 1-hour or 24-hour time points, indicating a sustained mechanism of resistance (**Fig 6F**). To further clarify the resistance mechanism of these lines to BRAFi, we quantified levels of GTP-bound RAS in parental and BRAFi-resistant lines (**Fig 6G**). Increased RAS activity was observed in two dabrafenib-resistant (DBTRG and NGT41) lines and the vemurafenib-resistant line (MAF794). We measured total ARAF and CRAF expression, as isoform switching is a recognized mechanism of resistance, and observed increased expression in some of the resistant lines (**Fig 6G**) (42). Increased phosphorylation of AKT and EGFR were also observed in both DBTRG and NGT41 lines.

**Figure 6.**
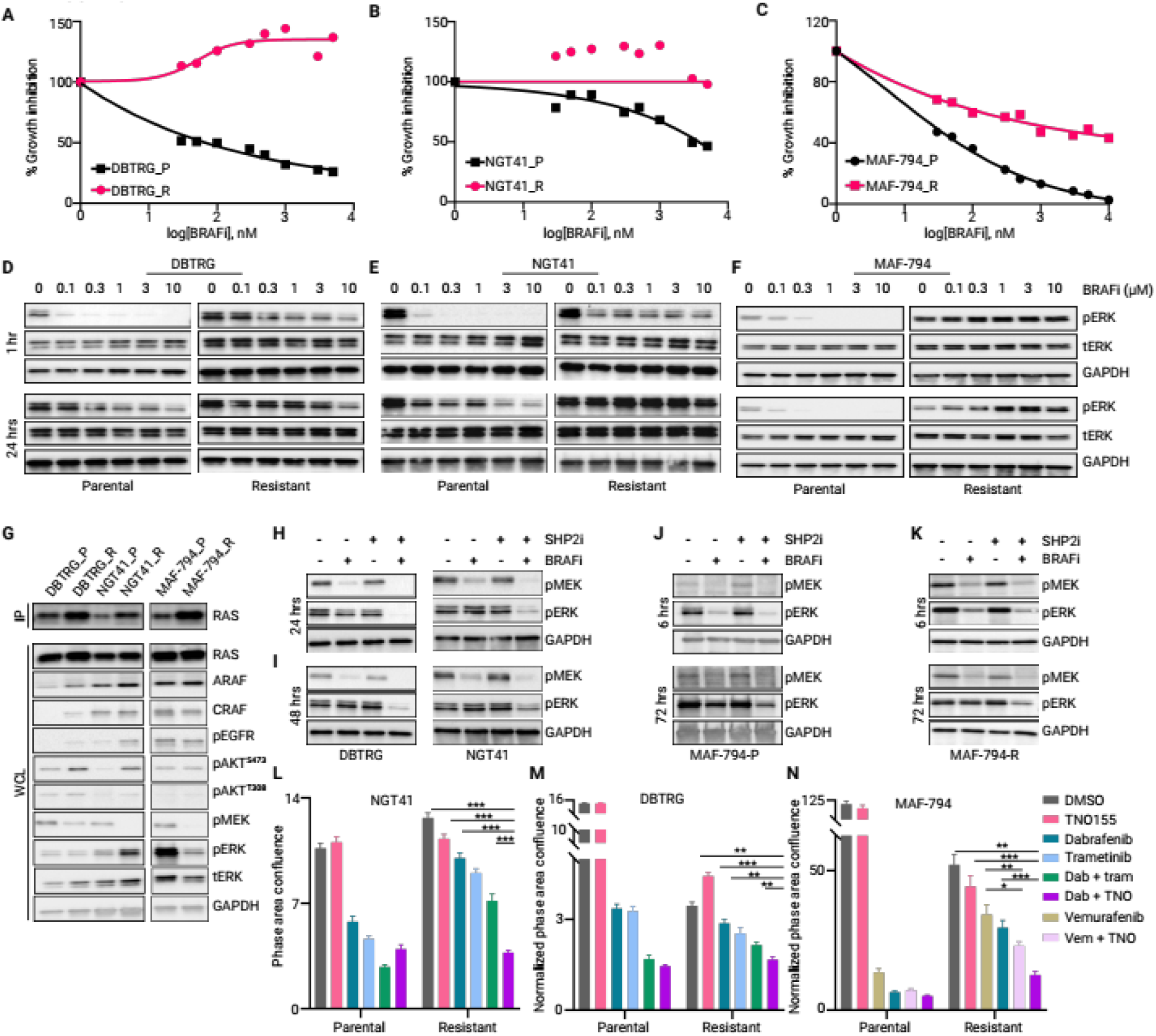
Combination of BRAFi and SHP2i overcomes BRAFi-resistance. A, B) Dose-response curves of parental and resistant lines treated with increasing doses of dabrafenib (0.1 µM-10 µM) or C) vemurafenib. D, E) Immunoblots evaluating ERK activity in response to dabrafenib or (F) vemurafenib monotherapy (100nM-10µM), at 1 and 24 hours post treatment. G) RAS-GTP pulldown assay in parental and resistant lines at baseline (24 hours after plating), IP=IP: RAF1-RBD, WCL= whole cell lysate. H, I) DBTRG and NGT41 dabrafenib-resistant lines were treated with DMSO, TNO155 (3 µM), dabrafenib (100 nM), or the combination for 24 or 48 hours then lysates were probed for the indicated proteins. Immunoblots of ERK activity in J) MAF794-P (parental) cells or K) MAF794-R cells resistant to vemurafenib (10 µM) following treatment with dabrafenib (100 nM), TNO155 (3 µM), the combination, then measured at 6 or 72 hrs. L, M) BRAFi-resistant lines, including dabrafenib-resistant NGT41 and DBTRG, and N) vemurafenib-resistant lines, along with parental controls, were treated with DMSO, BRAFi (300 nM dabrafenib), SHP2i (3 µM TNO155), or the combination for four days. Cell growth was monitored using Incucyte and normalized to time 0. Error bars indicate mean ± SEM (*p ≤0.05, **p ≤0.001, ***p ≤0.0001).

We then evaluated whether dabrafenib-resistant lines could be sensitized to BRAFi by the addition of a SHP2i. We observed the successful inhibition of pERK with combination therapy at 24 and 48 hours following the addition of SHP2i in some lines, and up to 72 hours in the vemurafenib-resistant cell line MAF794, possibly reflective of the longer half-life of vemurafenib (**Fig 6H-K**) (43). The combination of BRAFi and SHP2i also resulted in significantly greater cell growth inhibition compared to either monotherapy in all BRAFi-resistant cell lines (**Fig 6L-N**). BRAFi combined with SHP2i was superior to BRAFi combined with MEKi in resistant lines as well (**Fig 6L-M**). Furthermore, in the vemurafenib resistant line, dabrafenib combined with TNO155 was superior to vemurafenib combined with TNO155 (**Fig 6N**). Together these results demonstrate the addition of a SHP2i can overcome acquired resistance to BRAFi.

### Combined inhibition demonstrates efficacy *in vivo*

To assess the *in vivo* effects of combined BRAFi with SHP2i, we utilized heterotopic (flank) xenografts from two *BRAF* V600E mutant glioma cell-line models (**Fig 7A,B**) and one patient derived xenograft (PDX, **Fig 7C**). Mice were administered dabrafenib, TNO155, the combination, or vehicle twice daily. Tumors in mice receiving the combination exhibited significant growth inhibition in all three models, which was sustained over the entire treatment period (**Fig 7A-C**, **S7A, B**). Not only did the PDX tumors fail to grow with combination therapy, but they shrank from baseline, underscoring the potency of the combination in this PDX (**Fig 7D, E**). Immunoblot analysis at the end of study demonstrated ongoing inhibition of pERK with the combination, but increased levels of pERK in the dabrafenib group relative to vehicle, consistent with compensatory feedback (**Fig 7F**).

**Figure 7:**
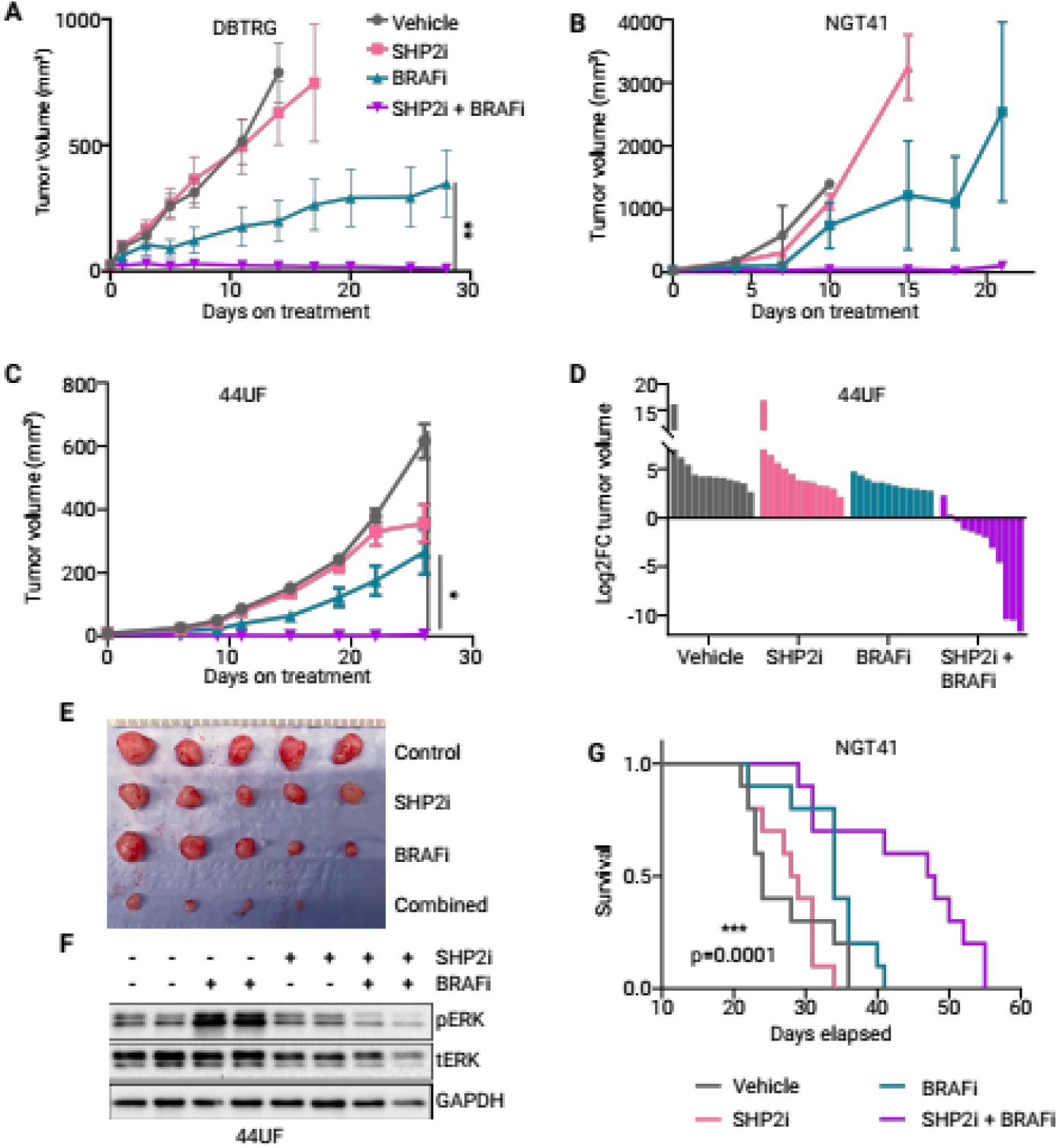
Combined small molecule inhibition demonstrates efficacy *in vivo*. A/B) Tumor volumes of heterotopically implanted BRAF-V600E cell lines (DBTRG-5MG and NGT41) treated with dabrafenib (15 mg/kg), TNO155 (7.5 mg/kg), or the combination twice daily (5 days on 2 days off) via oral gavage for up to 28 days. C) Tumor volumes and D) waterfall plot depicting log_2_ fold change in a patient-derived xenograft (PDX 44UF) heterotopic model, treated as above. Each bar represents an individual tumor. E) Representative image of tumors from five mice in each cohort of 44UF heterotopic xenografts. F) Immunoblots from PDX xenografts harvested at the end of treatment were probed for pERK relative to total ERK and GAPDH. G) Orthotopic, intracranially-implanted NGT41 xenografts were treated as above for 5 weeks starting from day 7 post-implantation. Survival was recorded by an independent, blinded evaluator. Hazard Ratio (log rank) of the combination versus dabrafenib was 2.514. Error bars represent mean ± SEM (*p ≤0.05, ***p ≤0.0001).

We next investigated the efficacy of dabrafenib with TNO155 in an intracranial tumor model. Mice were implanted intracranially with NGT41 and treated beginning five days post-transplant until removal from study (6 days on, 1 day off). The primary endpoint of survival was assessed by an independent, blinded evaluator. Mice treated with the combination lived significantly longer than controls or either monotherapy (p = 0.0097, Log Rank test) with a hazard ratio of 2.514 between dabrafenib-treated and combination-treated cohorts (**Fig 7G**). The mice treated with the combination of dabrafenib and TNO155 displayed no signs of adverse effects on body weight or grooming throughout the study (**Fig S7C-D**). Furthermore, higher doses of either monotherapy did not yield better effects on tumor volume inhibition (**Fig S7E**).

## Discussion

Adaptive and acquired resistance pose significant hurdles to the effectiveness of targeted therapy. The recent successes of BRAF/MEK combination therapy in glioma and the paucity of other effective therapies have shifted attention to understanding and overcoming therapeutic resistance. Here, we show that adaptive resistance in glioma often occurs through reactivation of MEK-ERK signaling and can be overcome by simultaneous inhibition of a critical signaling node within the same pathway – SHP2. Combined SHP2-BRAF inhibition suppresses ERK activity more deeply and durably than either monotherapy. Moreover, the combination overcomes resistance to BRAFi in some cell lines and demonstrates *in vivo* efficacy and tolerability.

Our data presented herein build upon human glioma data that demonstrate treatment-emergent resistance often occurs through acquired mutations that maintain ERK dependence (9, 11). In those studies, a large subset of gliomas did not have an identified resistance mechanism. Here we show that ERK signaling reactivation is often present at time of treatment-emergent resistance. Moreover, this reactivation is consistent with the pattern of RAS activity seen in *BRAF* V600E glioma lines upon adaptive resistance, underscoring the significance of RAS-ERK signaling for survival and proliferation in these gliomas.

While resistance to targeted therapy often emerges through reactivation of ERK signaling, the selection of a next-line therapeutic strategy requires a careful examination of genomic and proteomic features that confer selective advantages and drive resistance. A broad selection of small-molecule inhibitors in the RAS-ERK pathway are being developed and have a range of different chemical properties and target proteins. Here, we selected SHP2 as a target for combination with BRAF-directed targeted therapy due to the central obligatory function of SHP2 for RAS signaling (18, 19). Moreover, SHP2 plays a key role in resistance to targeted therapy in other *BRAF* mutant cancers (21, 44–47), and combined pathway inhibition shows increased potency in ERK pathway dysregulated cancers (21, 45–47). Our work demonstrates that, in glioma, SHP2 co-inhibition can extend the durability of response to a BRAFi in *BRAF* mutant glioma models both sensitive or resistant to BRAFi, suggesting that combined vertical inhibition of ERK signaling may have therapeutic potential. While we demonstrated that synergy occurs through inhibition of compensatory RAS activation, this model did not take into account the role of autophagy in resistance (10), nor the potential role of the tumor microenvironment in modulating sensitivity (48).

Targeted therapy combinations for gliomas require blood-brain barrier penetration. Dabrafenib and trametinib each have demonstrated efficacy as monotherapy in low-grade glioma, indicative of blood-brain barrier penetration (4, 49). TNO155 is an allosteric inhibitor that, when bound, modifies the conformation of SHP2, stabilizing its auto-inhibited state (50). While it is currently being tested alone or in combination with other small molecule inhibitors in clinical trials (NCT03114319), there are no data for its brain penetration, though its precursor compound, SHP099, has shown brain penetration and efficacy in orthotopic glioblastoma models (51, 52). Our data support modest efficacy in an orthotopic glioblastoma model despite good activity in the same cell line implanted heterotopically. This finding may indicate reduced blood-brain barrier penetration and additional studies will need to be performed to optimize pharmacokinetics for clinical translation. Other allosteric SHP2 inhibitors are also in clinical development, some with potential for blood-brain barrier penetration (53).

Balancing tolerability with efficacy can be a significant challenge when combining small molecule targeted inhibitors. Profound inhibition of MEK-ERK signaling can lead to significant toxicity through ERK suppression in normal cells. For example, while combined SHP2 and MEK inhibition has shown efficacy in some cancer models, some toxicity was observed in human participants (52, 54). The combination of BRAFi and SHP2i evaluated here, however, shows promise for improved tolerability due to the specificity of dabrafenib for *BRAF* V600E, potentially improving the therapeutic index. Similar to the combination of BRAFi plus MEKi, BRAFi plus SHP2i should result in an improved toxicity profile due to paradoxical activation from the BRAFi, leading to opposite effects on ERK phosphorylation in normal cells (55). Here we found the combination of dabrafenib with TNO155 was active in *BRAF* V600E mutant *in vivo* and well-tolerated by mice.

Overall, our study elucidates the functional role of SHP2 in adaptive resistance to BRAFi in gliomas and highlights the therapeutic potential of combined BRAF and SHP2 inhibition for preventing or overcoming resistance. This combination demonstrates potent anti-tumor efficacy *in vitro* and *in vivo*, providing a promising strategy for overcoming resistance and improving outcomes in patients with *BRAF* V600E mutant gliomas. Further investigation is warranted to validate these findings and translate them into clinical practice.

## Authors’ contributions

A. A. Ayanlaja: Investigation, visualization, validation, project administration, methodology, writing– original draft, writing–review and editing. M. Chang: Formal analysis, validation, visualization, writing– review. M. Ioannou: Investigation, methodology. K. Lalwani: Investigation, methodology, and analysis. S. J. Wang: methodology, writing–review and editing. Jagtap: Investigation, methodology. Y. Yang: methodology, Formal analysis. R. D. Gartrell: Formal analysis and validation, resources, writing–review and editing. C. Pratilas: resources, supervision, writing–review and editing. K.C. Schreck: Conceptualization, resources, formal analysis, supervision, funding acquisition, writing–original draft, project administration, writing–review and editing.

## Acknowledgements

This work was funded by DOD grant (W81WH2110251), Doris Duke Early Clinician Investigator Award, AACR-PLGA (20-60-63-SCHR), Cigarette Restitution Fund, the Matthew Larson Foundation, and SKCCC Core grant (UM1-CA-137443). The Gartrell Lab (R.D.G and Y.Y) is supported by Rally Foundation, Swim Across America, and Children’s Cancer Foundation. TNO155 was provided by Novartis Institute for Biomedical Research (NIBR).

