## Supplementary figures for "Combined inhibition of SHP2 overcomes adaptive resistance to type 1 BRAF inhibitors in BRAF V600E-driven high-grade glioma"

**SUPPLEMENTARY FIGURE 1**

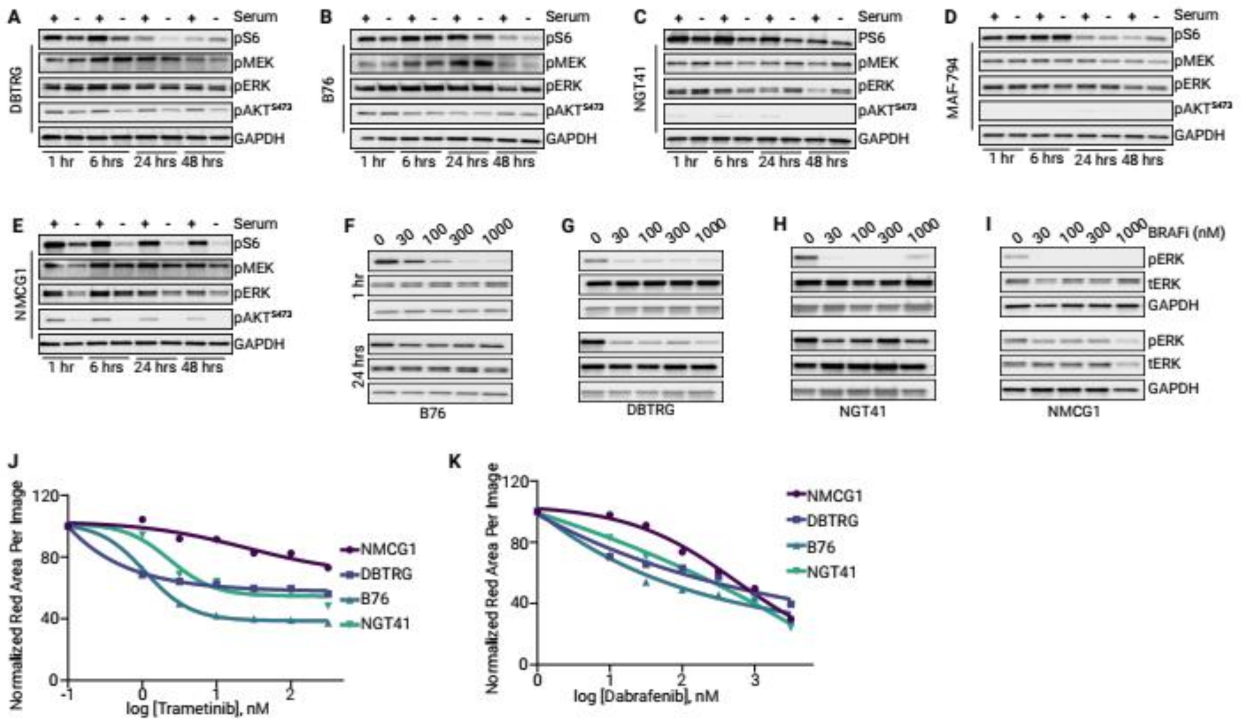


**Supplementary Figure 1:** Administration of 10-15% fetal bovine serum compared with serum starvation over time in A) DBTRG, B) B76, C) NGT41, D) MAF-794, and E) NMCG1 glioma cell over time. F) Immunoblots from cells treated with increasing doses of dabrafenib (30nM-1000nM) showing ERK signaling at 1 and 24 hours in B76, G) DBTRG, H) NGT41, I) and NMCG1 cells. Dose-response curves to J) trametinib or K) dabrafenib monotherapy in *BRAF* V600E mutant glioma cells treated with increasing drug doses for five days.

SUPPLEMENTARY FIGURE 2

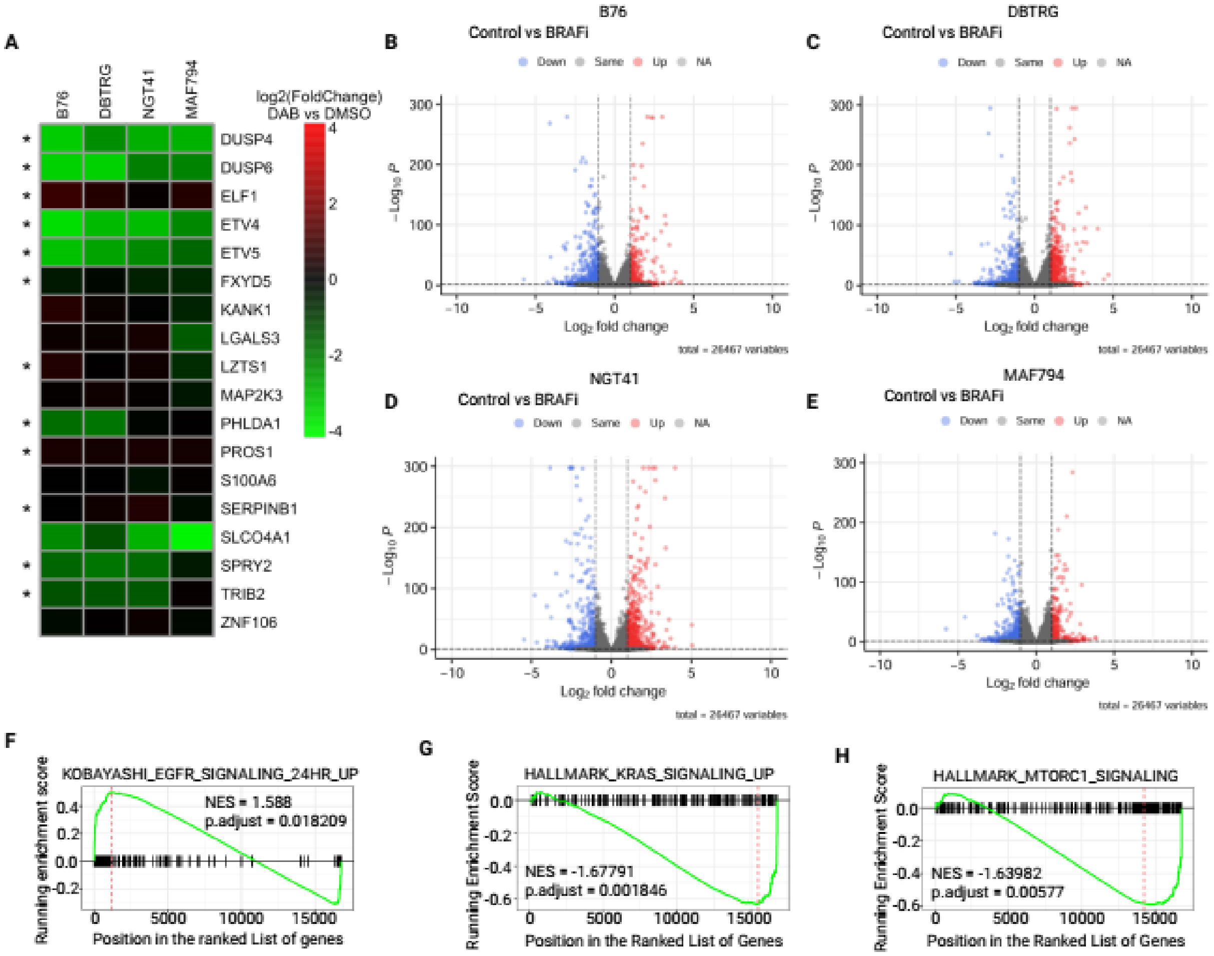


**Supplementary Figure 2:** A) Expression of an 18-gene signature for tumor-independent MEK functional output showing $log_2FC$ with respective p values of transcripts in four cell lines treated with BRAFi vs control. Volcano plots illustrating positive and negative LFC (log fold change) transcripts in BRAFi-treated (dabrafenib, 100nM)-treated B) B76, C) DBTRG, D) NGT41), and E) MAF794 cells compared to controls. F-H) Barcode plots of selected significantly altered pathways relevant for MAPK signaling following BRAFi compared to vehicle.

SUPPLEMENTARY FIGURE 3

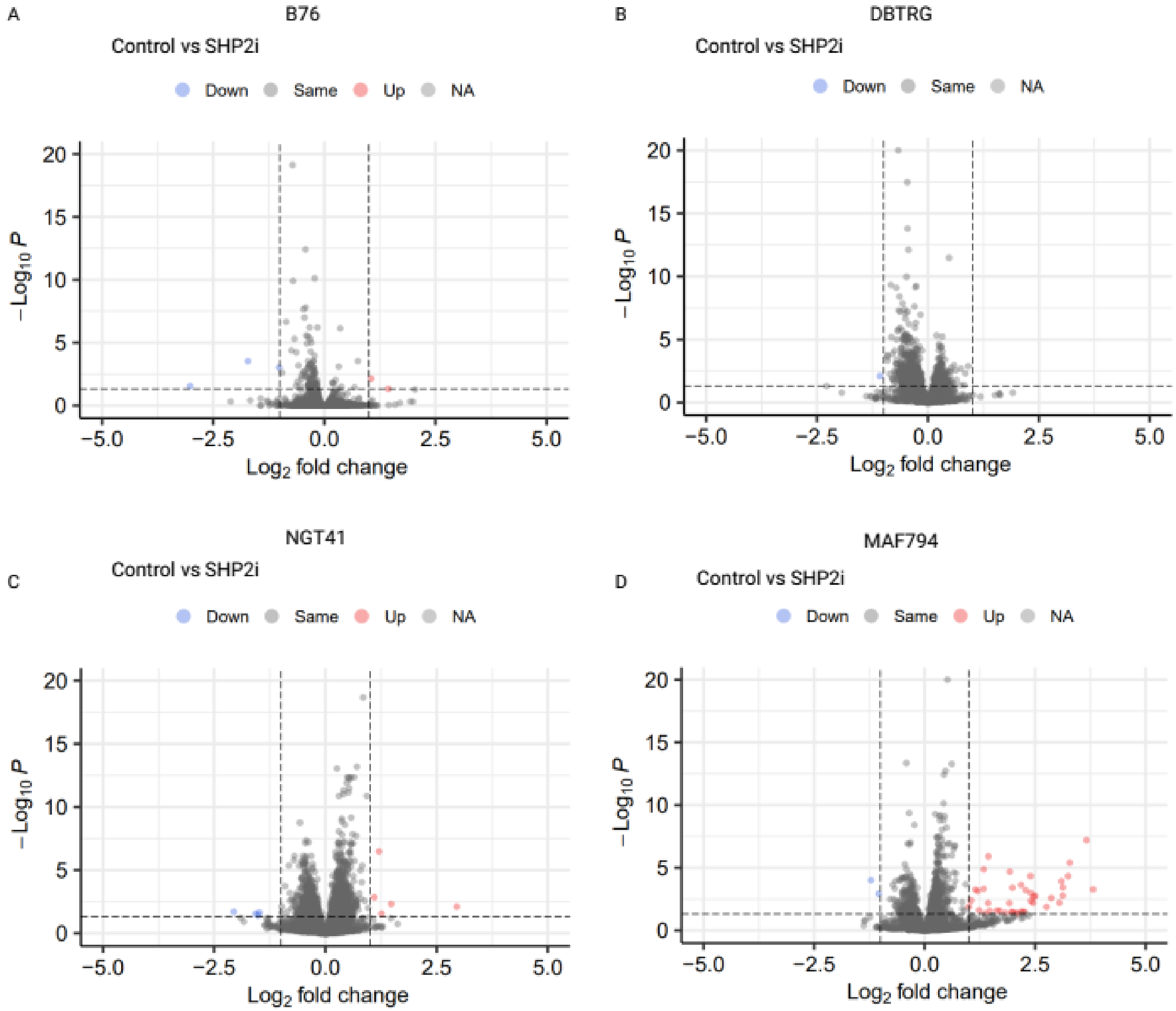


**Supplementary Figure 3:** Volcano plots illustrated positive and negative LFC (log fold change) transcripts in SHP2i-treated (TNO155, 3 µM)-treated A) B76, B) DBTRG, C) NGT41), and D) MAF794 as compared to vehicle.

**SUPPLEMENTARY FIGURE 4**

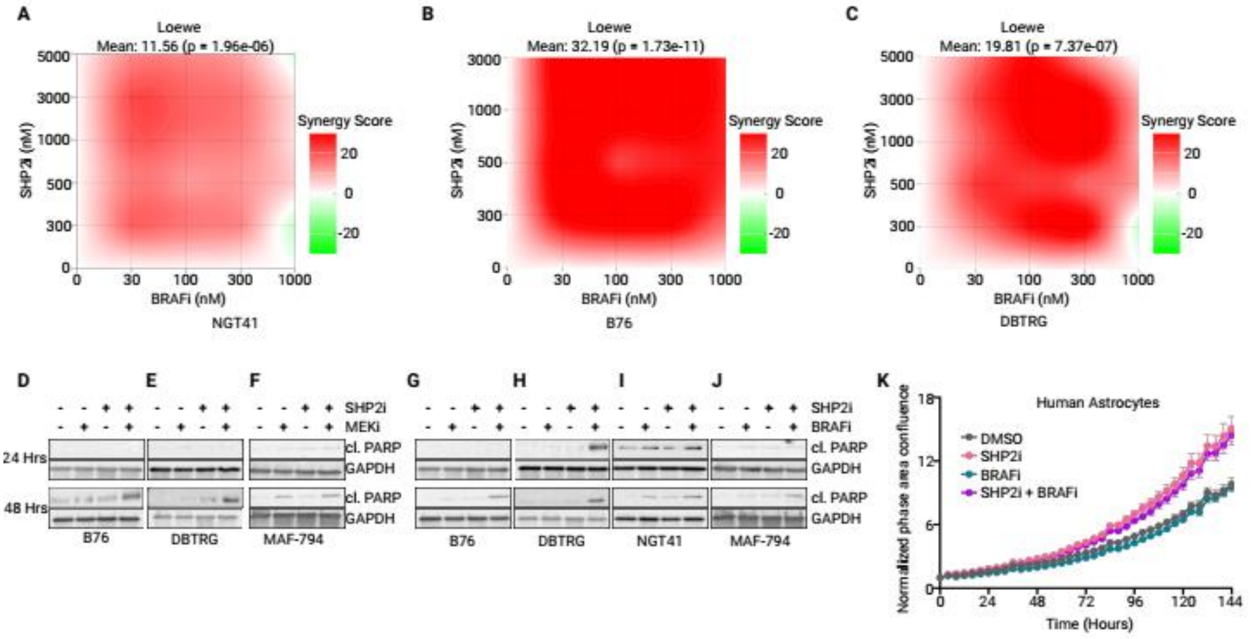


**Supplementary Figure 4:** A-C) Loewe's synergy heatmaps of TNO155 combined with dabrafenib in NGT41, B76, and DBTRG cells. D-F) Immunoblots of cleaved PARP expression in B76, DBTRG, and MAF-794 lines at 24 and 48 hours after treatment with MEKi (30nM, trametinib), SHP2i (3 µM, TNO155) or the combination. Samples are from the same experiment as main Figure 3C. G-J) Immunoblots of cleaved PARP expression in B76, DBTRG, NGT41, and MAF-794 lines at 24 and 48 hours after treatment with BRAFi (100 nM, dabrafenib), SHP2i (3 µM, TNO155), or the combination. Samples are from same as experiment as main Figures 4E or 6J. H) Growth of human astrocytes measured with Incucyte live cell-imager after treatment with dabrafenib (100 nM), TNO155 (3 µM), or the combination.

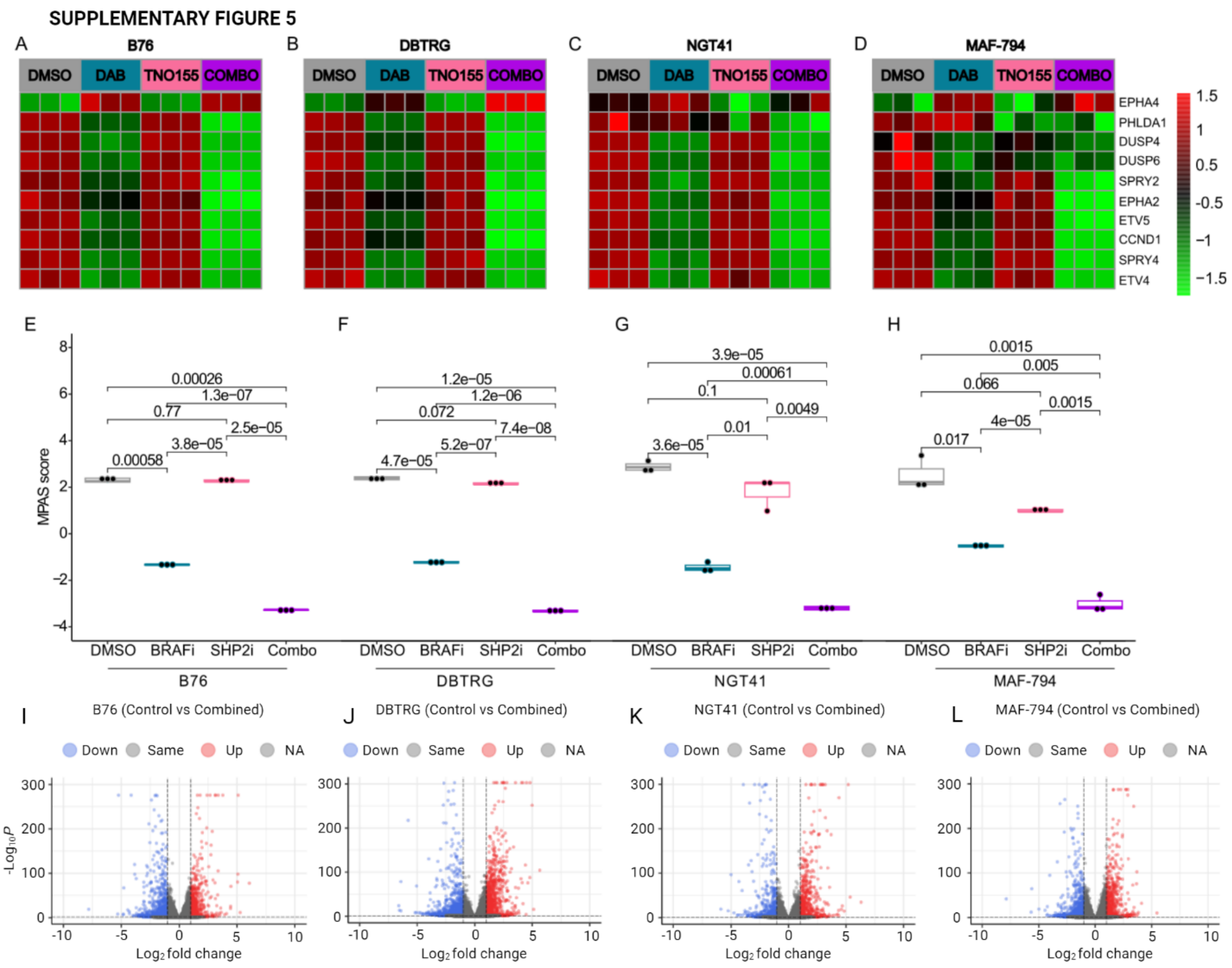


**Supplementary Figure 5.** Heatmap of $\log_2$ TPM of the 10 genes in the MPAS signature in triplicates for A) B76, B) DBTRG, C) NGT41 and, D) MAF-794 cells treated with DMSO, dabrafenib, TNO155, or the combination. E-H) Box-and-whisker plots displaying mean scaled expression of MPAS score for B76, DBTRG, NGT41, and MAF794 cell lines treated with DMSO, dabrafenib, TNO155, or combination. I-L) Volcano plots showing positive and negative LFC (log fold change) transcripts in BRAFi + SHP2i (dabrafenib-100 nM, TNO155-3 μM)-treated A) B76, B) DBTRG, C) NGT41), and D) MAF794 as compared to controls.

**SUPPLEMENTARY FIGURE 7**

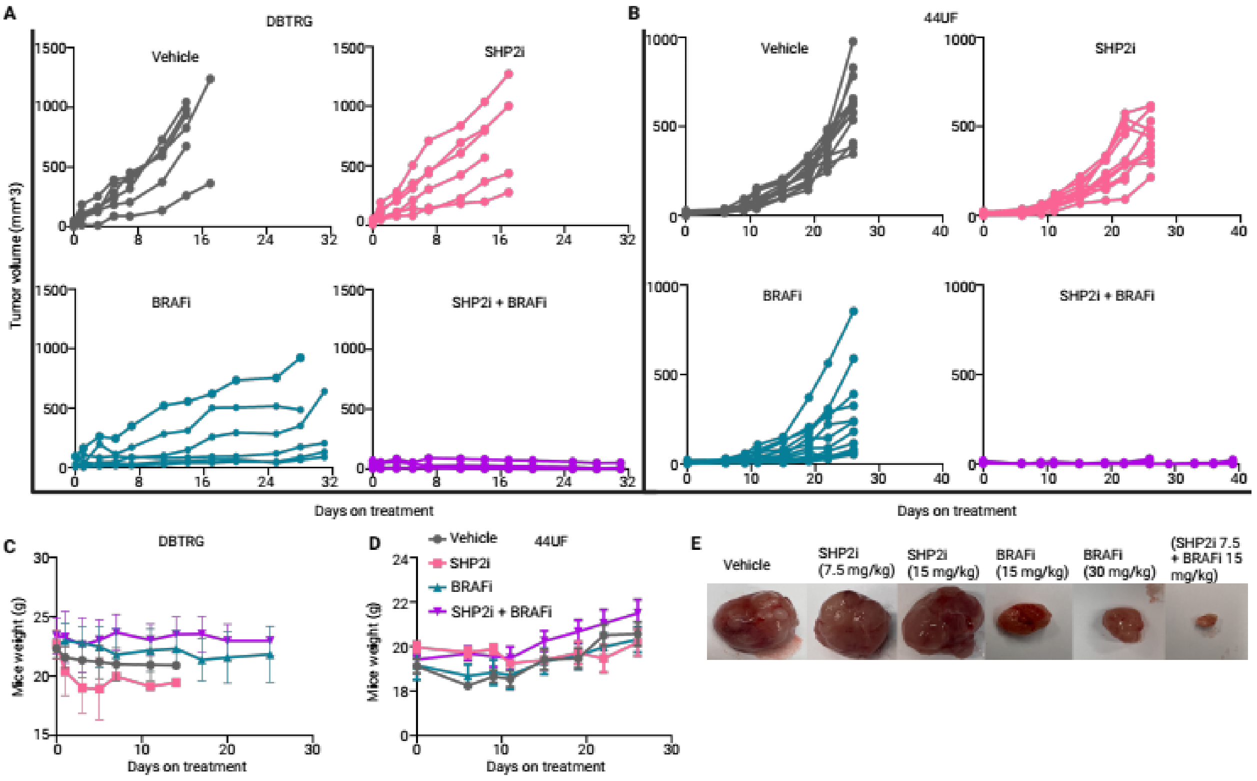


**Supplementary Figure 7:** A/B) spider plot of individual tumor volume in mice flanks for DBTRG and 44UF experiments. C/D). Mice weight over time on treatment for each study. E) Tumor size at end of study for higher monotherapy doses of TNO155 and dabrafenib in NGT41 xenografts.
